# Stem cell homeostasis regulated by hierarchy and neutral competition

**DOI:** 10.1101/2022.03.23.485259

**Authors:** Asahi Nakamuta, Kana Yoshido, Honda Naoki

**Affiliations:** Laboratory of Theoretical Biology, Graduate School of Biostudies, Kyoto University, Yoshidakonoecho, Sakyo, Kyoto, 606-8315, Japan; Faculty of Science, Kyoto University, Yoshidakonoecho, Sakyo, Kyoto, 606-8315, Japan; Laboratory of Data-driven Biology, Graduate School of Integrated Sciences for Life, Hiroshima University, Kagamiyama, Higashi-hiroshima, Hiroshima, 739-8526, Japan; Theoretical Biology Research Group, Exploratory Research Center on Life and Living Systems (ExCELLS), National Institutes of Natural Sciences, Okazaki, Aichi, 444-8787, Japan

## Abstract

Tissue stem cells maintain themselves through self-renewal while constantly supplying differentiating cells. As a mechanism of stem cell homeostasis, two distinct models were proposed: the classical model states that there are hierarchy among stem cells and master stem cells provide stem cells by asymmetric divisions, whereas the recent model states that stem cells are equipotent and neutrally competing each other. However, its mechanism is still controversial in several tissues and species. Here, we developed a mathematical model linking these two models. Our theoretical analysis showed that the model with the hierarchy and neutral competition, called the hierarchical neutral competition (hNC) model, exhibited bursts in clonal expansion, unlike existing models. Furthermore, the scaling law in clone size distribution, thought to be a unique characteristic of the recent model, was satisfied in the hNC model. Based on these findings, we proposed a criterion for distinguishing the three models by experiments.

## Introduction

All cells in our body are derived from stem cells, which maintain their population through self-renewal while constantly producing differentiated cells to achieve homeostasis in any tissue. This property of tissue stem cells is vitally important for homeostasis in organisms, and defects in stem cell homeostasis cause various diseases such as cancer and infertility^1–3^. In the stem cell homeostasis, fate asymmetry is achieved, in which half of the daughter cells from the entire stem cell population is retained as stem cells and the other half is directed toward differentiation. To date, two distinct models have been suggested to explain the mechanism of stem cell homeostasis, i.e., how the fate asymmetry is achieved^4,5^: the hierarchical model and neutral competition (NC) model. However, the mechanism of stem cell homeostasis remains controversial in many species and tissues.

The hierarchical model has traditionally been accepted. In this model, there are two hierarchies of stem cells: master and non-master stem cells (**Fig. 1a**). Master stem cells are the most undifferentiated stem cell population consisting of a limited number of stem cells, while non-master stem cells are more differentiated and irreversibly directed toward differentiation. For stem cell homeostasis, one master stem cell generates one master stem cell and one non-master stem cell through asymmetric division (**Fig. 1b**). This asymmetry at the single-cell level achieves fate asymmetry in the entire stem cell population, where loss of stem cells by their differentiation is compensated by asymmetric division of master stem cells, ensuring maintenance of the stem cell population. In *Drosophila*, stem cells in the germline and developing central nervous system have been shown to undergo invariant asymmetric cell divisions, giving rise to one stem cell and one differentiating cell^4,6^, strongly supporting the hierarchical model. This model has also been widely considered in mammalian stem cell homeostasis. In classical models of epidermal and spermatogenic stem cells, a limited number of stem cells are thought to maintain each tissue homeostasis^7–9^.

**Fig. 1.**
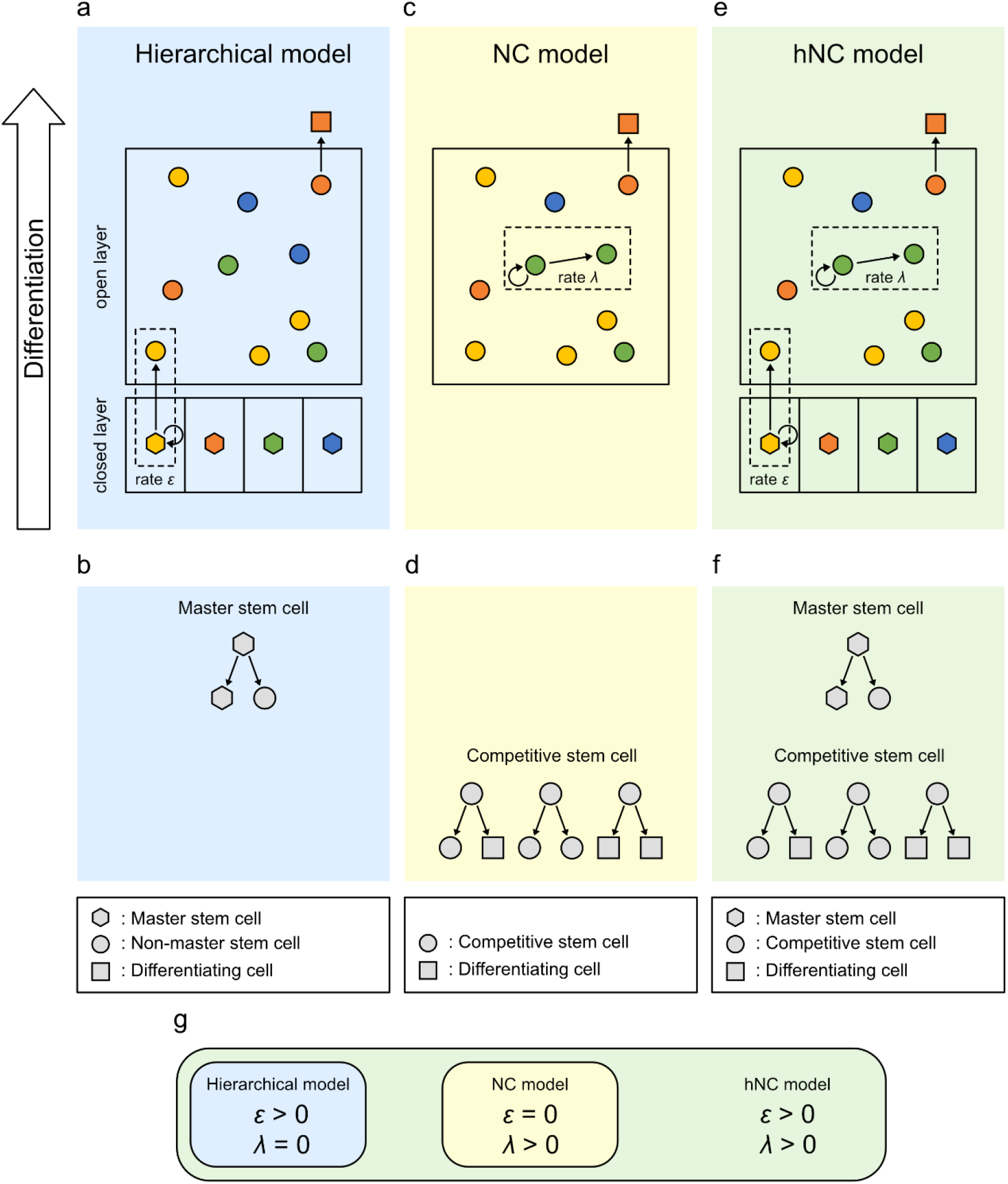
Three distinct biological models for stem cell homeostasis. (a) The hierarchical model. Loss of stem cells is compensated by the asymmetric division of master stem cells (hexagon) in the closed layer. In the open layer, non-master stem cells (circles) do not compete with each other and are directed to differentiation. Each color indicates each clone derived from the common master stem cell. (b) Diagram of patterns of stem cell divisions in the hierarchical model. (c) The NC model. Loss of stem cells is compensated by the proliferation of competitive stem cells (circles) in the open layer. Each color indicates each clone derived from the common competitive stem cell. (d) Diagram of patterns of stem cell divisions in the NC model. (e) The hierarchical neutral competition (hNC) model. Loss of stem cells is compensated by the asymmetric division of master stem cells (hexagon) in the closed layer and proliferation of competitive stem cells (circles) in the open layer. Each color indicates each clone derived from the common master stem cell. (f) Diagram of patterns of stem cell divisions in the hNC model. (g) Relationships between the three models. The mathematical model seamlessly represented all three biological models depending on two parameters, *ε* and *λ*, which represent the proliferation rates of master stem cells and competitive stem cells, respectively.

Another distinct model, known as the neutral drift model or stochastic model, has recently attracted attention^10^. We refer to this model as the NC model to focus on the clonal expansion process occurring through neutral cell-cell competition. In the NC model, all stem cells are equipotent without the hierarchy among stem cells assumed in the hierarchical model, we call all these stem cells “competitive stem cells” (**Fig. 1c**). This model assumes that the fate of each stem cell is not fixed: a stem cell generates two stem cells or two differentiating cells by symmetric division or one stem cell and one differentiating cell by asymmetric division (**Fig. 1d**). Therefore, fate asymmetry is achieved at the cell population level rather than at the single-cell level, and loss of one stem cell is compensated by the symmetric division of the other stem cell into two stem cells. Thus, stem cells neutrally compete with each other, leading to neutral drift in clonal expansion. The NC model was originally suggested based on the observation of the random fate of stem cells^11,12^ and several inducible genetic labeling studies^13,14^. Recently, Klein and Simons performed mathematical analysis of the NC model and showed that stem cell clonal expansion follows the scaling law, in which the scaled probability distribution of the stem cell population size of each clone is universal over time^5^. Several experimental studies involving long-term lineage tracing of stem cell clones confirmed this scaling law in mouse epidermal, intestinal, and spermatogenic tissues, supporting the NC model as the mechanism of stem cell homeostasis^15–17^. Therefore, the NC model has recently been regarded as another candidate for the mechanism of stem cell homeostasis in mammalian tissues.

However, a phenomenon that cannot be predicted by either existing model alone has been reported. A study using genetic barcoding of spermatogonial stem cells in mice reported that offspring were derived non-randomly from all clonal types of spermatogonial stem cells, and that offspring of specific clones were periodically observed, suggesting that the population size of each stem cell clone exhibited transient burst-like patterns repeatedly^18^. In the hierarchical model, the clone population size is stably maintained because each master stem cell compensates for the loss of stem cells without competition, whereas in the NC model, a few clones dominate the population and other clones end up in extinction owing to the neutral evolution of the stem cell clones. Because both the hierarchical model and the NC model are not able to explain burst-like dynamics of stem cell clones, there should be a missing factor for understanding stem cell homeostasis.

To address this missing factor and the controversy in the mechanism of mammalian stem cell homeostasis, we focused on the fact that two existing models are not necessarily conflicting. In fact, both models have been supported by data from different experimental systems, leading to different results. Thus, the same phenomenon has been possibly observed from the different aspects. Based on this idea, in this study, we presented a new and comprehensive mathematical model that seamlessly links two existing models. In addition, this mathematical model can represent the intermediate model we named hierarchical neutral competition (hNC) model, in which both the supply of stem cells by master stem cells and competition between stem cell clones are compatible. Through numerical simulation and mathematical analysis, we revealed that the hNC model exhibited burst-like patterns in the clone dynamics of stem cells. Furthermore, we showed that the scaling law in clone size distribution, which was proposed as the indicator of the NC model, was satisfied not only in the NC model but also in the hNC model. Based on these findings, we proposed a possible criterion for distinguishing these three conditions experimentally.

## Results

### General model for stem cell homeostasis

To examine the effect of the presence of master stem cells and/or neutral competition among stem cells on clonal dynamics in stem cell homeostasis, we developed a mathematical model that comprehensively represents the hierarchical, NC, and their intermediate model named as the hNC models (**Fig. 1a–f**). The hNC model has two hierarchies of stem cells as in the hierarchical model: master stem cells in the closed layer and competitive stem cells in the open layer, while it also has neutral competition among competitive stem cells as in the NC model (**Fig. 1e**). In the closed layer, master stem cells supply competitive stem cells to the open layer while undergoing self-renewal via invariant asymmetric divisions. In the open layer, competitive stem cells compete with each other while producing differentiating cells; when a competitive stem cell is lost through differentiation or apoptosis, it is compensated either through the supply from master stem cells or symmetric division of other competitive stem cells at rates of *ε* and *λ*, respectively (**Fig. 1f**).

In the simulation of our model, at each step, one competitive stem cell was randomly selected for differentiation and excluded from the open layer. Next, a master stem cell or competitive stem cell was selected with a weighted probability defined by the parameters *ε* and *λ*, which supplies one competitive stem cell to maintain the total number of stem cells. From a theoretical perspective, this model was extended from the Moran process^19^, which generally describes the evolutionary population dynamics of two species, by introducing multiple species and an external supply from master populations outside of competition.

This mathematical model can represent three biological models by changing parameters *ε* and *λ* (**Fig. 1g**). The condition under which competitive stem cells do not proliferate (*λ* = 0), indicating the absence of neutral competition, corresponds to the hierarchical model. In contrast, the condition under which master stem cells do not supply competitive stem cells (*ε* = 0), indicating the absence of master stem cells, corresponds to the NC model. Additionally, the mathematical model represents the intermediate condition between the hierarchical and NC models named as the hNC model, that is *λ* > 0 and *ε* > 0.

### Clonal bursts were generated in the hNC model

To investigate the dynamics of the three different biological models (hierarchical, NC, and hNC models) for explaining stem cell homeostasis, we performed numerical simulations of our mathematical model using different values for the proliferation rate of master stem cells *ε* and that of competitive stem cells *λ*. In the hierarchical model (*ε* > 0, *λ* = 0), no clone dominated the open layer, and the populations of all clones fluctuated around the averages because master stem cells continuously compensated for the loss of stem cells from the closed layer (**Fig. 2a**). In contrast, in the NC model (*ε* = 0, *λ* > 0), only one clone dominated the open layer, and all other clones were extinguished through neutral, random drift of clonal populations (**Fig. 2b**). Interestingly, in the hNC model (0 < *ε* ≤ *λ*), each clone population showed intermittent and repeated expansion and contraction, like the bursts (**Fig. 2c**). These clonal bursts were generated by the mechanism that each clone headed to domination or extinction by neutral drift, but these two extreme results were not eventually achieved because of the infrequent supply from master stem cells, whose proliferation rate was lower than that of competitive stem cells.

**Figure 2.**
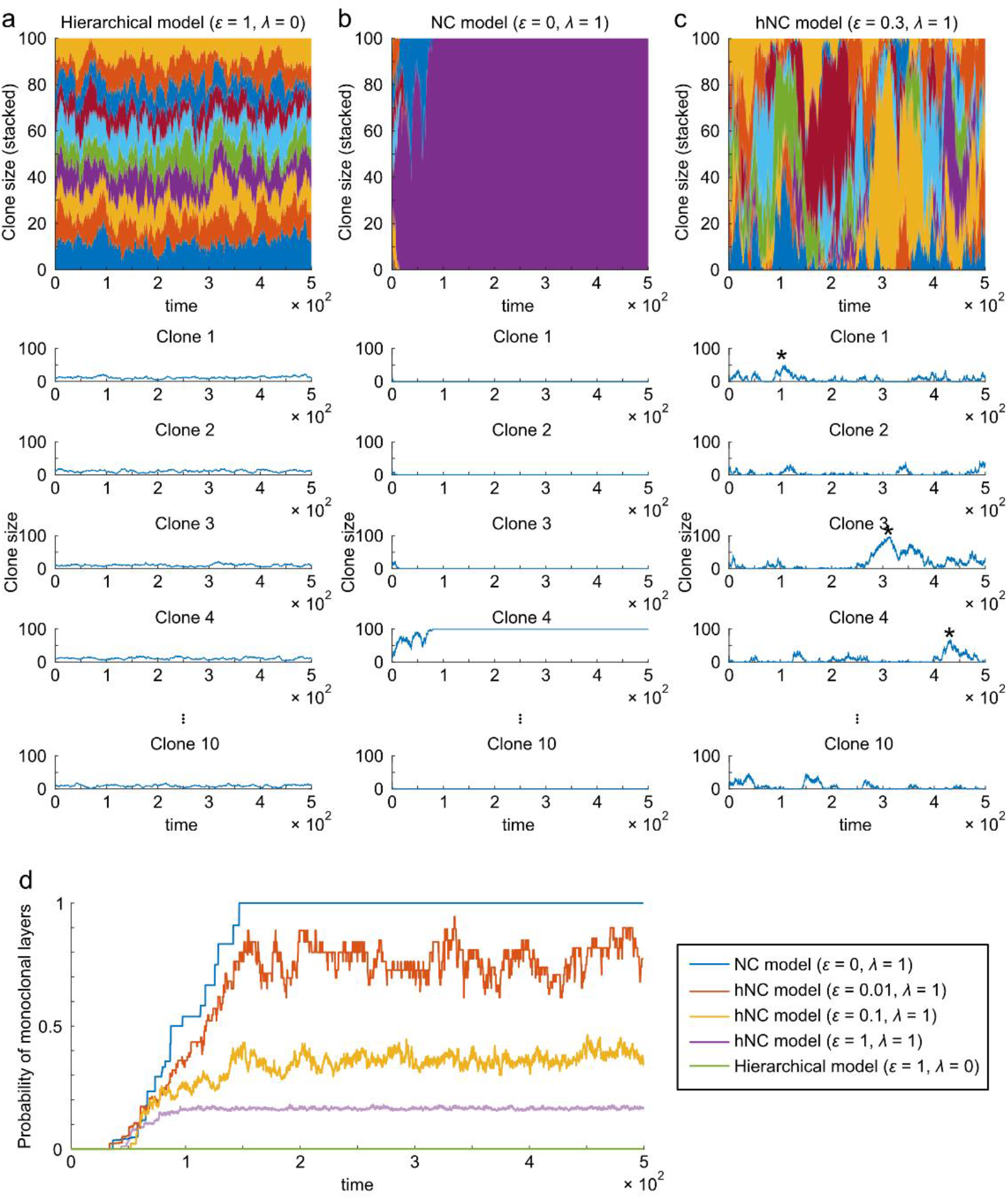
Clonal expansion in the hierarchical, NC, and hNC model. (a–c) Time-series of clone sizes with different proliferation rates of master stem cells (*ε*) and competitive stem cells (*λ*). In the simulation, there were 10 types of clones in 100 competitive stem cells in the open layer, in which the clone size of every clone was initially uniform, i.e., *n*_*k*_ = 10 at *t* = 0. Asterisks indicate representative bursts. In the top panels, fraction of each clone was displayed by different color. In the lower panels, the results of five representative clones within 10 clones are displayed. (d) The probability of monoclonal conversion in open layers over time. We repeated simulations 1000 times, and the probability that the open layer became monoclonal at each timepoint was displayed.

As another characteristic of clonal expansion, we examined monoclonal conversion, a phenomenon in which only one clone dominates the stem cell population. Using simulation, we quantified the probability of monoclonal conversion in the three models (**Fig. 2d**). Monoclonal conversion did not occur in the hierarchical model, whereas in the NC model, monoclonal conversion occurred; that is, the probability of monoclonal conversion increased over time and eventually converged to one in the NC model. In contrast, in the hNC model, the probability of monoclonal conversion increased over time and eventually reached and fluctuated around certain constants, approaching a larger value with a lower proliferation rate of master stem cells *ε*.

### Clonal bursts were caused by low proliferation of master stem cells

We mathematically analyzed how clonal bursts were generated as shown in Fig. 2 based on the theory of stochastic processes. We first derived the stationary probability distribution of the clone size (see **Methods**). We found that a smaller proliferation rate of master stem cells *ε* was associated with a higher probability of a large clone size (**Fig. 3a**), implying the existence of higher clonal bursts. Next, we directly analyzed the properties of clonal bursts. We defined a clonal burst (**Fig. 3b**) and derived the probability that a clonal burst of a certain height is generated and the expected duration of a clonal burst at a certain height (see **Methods**). By this theoretical analysis, we found that a lower proliferation rate of master stem cells increased and decreased the probabilities of larger and smaller bursts, respectively (**Fig. 3c** and simulation results in **Supplementary Fig. 1**). In addition, we found that a lower proliferation rate of master stem cells increased and decreased the durations of larger and smaller bursts, respectively (**Fig. 3d** and simulation results in **Supplementary Fig. 1**). These mathematical analyses were validated by confirming that the analytical solutions were reproduced in numerical simulations (**Fig. 3a, c, d**). Taken together, the proliferation rate of master stem cells relative to that of competitive stem cells dominantly determines the height and duration of stem cell clonal bursts.

**Figure 3.**
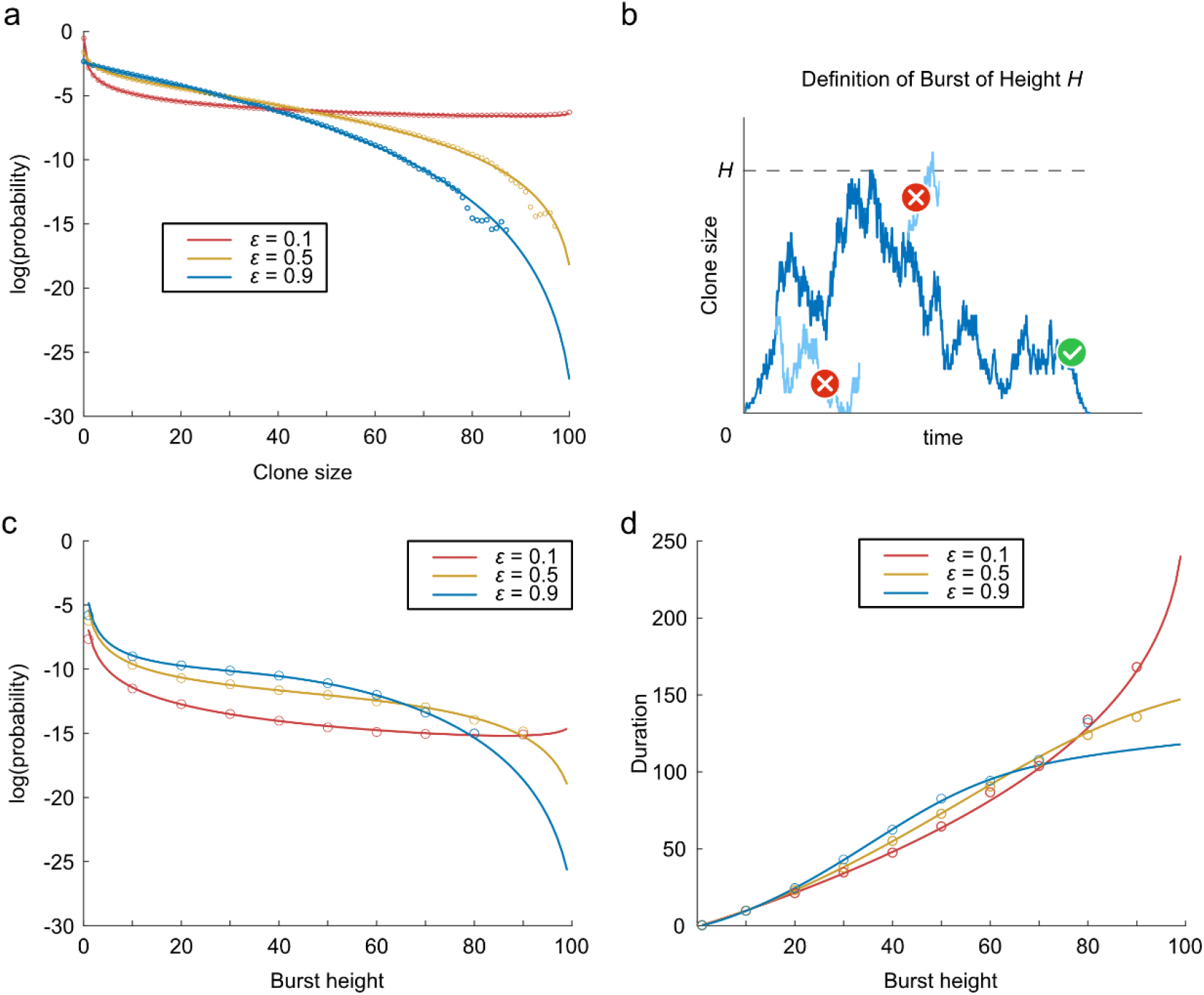
Clone size distribution and clonal bursts depending on the proliferation rate of master stem cells in the hNC model. (a) Stationary probability distribution of clone size with different proliferation rates of master stem cells *ε*. The solid line and dots indicate the probability distributions derived through mathematical analysis and numerical simulation, respectively. (b) Definition of clonal bursts. A burst of height *H* is defined as the dynamics in which the clone size changes from 0 to *H* without returning to 0, and then returns to 0 without reaching *H* + 1. (c) Probability of burst generation of each height *H* starting at clone size 0 depending on the proliferation rates of master stem cells *ε*. Solid lines indicate the analytical solution of the probability (equation (14)). Dots indicate the probability calculated by numerical simulation. (d) Expected duration of a clonal burst of each height *H* starting at clone size 0 depending on the proliferation rates of master stem cells *ε*. Solid lines indicate the analytical solution of the expected duration (equation (21)). Dots indicate the average duration calculated by the numerical simulation.

### Scaling law is satisfied in both the NC and hNC models

The existence of clonal bursts should be the indicator of the hNC model, consisting of hierarchy and neutral competition among stem cells, because the existing hierarchical and NC models cannot generate a clonal burst. On the other hand, Klein and Simons theoretically suggested that the NC model can be distinguished from the hierarchical model using the scaling law of the clone size distribution, in which the probability distribution of the stem cell population size of each clone, *Pn*(*t*), time-independently obeys the universal scaled distribution ^5^:

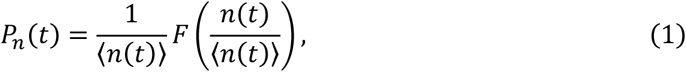

where *n*(*t*) and ⟨*n*(*t*)⟩ denote the clone size and its average at time *t*, respectively, and *F*(*x*) indicates the time-independent universal function, which was determined from the spatial dimension of the tissue of interest. In our situation in the NC model, *F*(*x*) was expected to obey an exponential distribution^20–22^. In fact, the dynamics of clonal expansion was experimentally monitored using lineage tracing of stem cells, which showed that the distribution of the clone size followed the scaling law in several tissues in mice^15,16,21^. Notably, the scaling law holds only in the early phase of clonal expansion and not in the late phase, during which all stem cell populations converge to almost monoclonal.

To investigate whether the clone size distribution satisfies the scaling law in the hNC model, we simulated our mathematical model with random labeling of one of the master and competitive stem cells. In the hierarchical model, the clone size distribution did not follow the scaling law (**Fig. 4a**). In contrast, the NC model obeyed a time-independent exponential distribution which followed the scaling law (**Fig. 4b**). These observations are consistent with previous findings. Furthermore, we observed scaling law of clone size distribution in the hNC model when the proliferation rate of the master stem cells was lower than or equal to that of the competitive stem cells (**Fig. 4c, d**), including the condition in which clonal bursts were generated, as shown in Fig. 2c and Supplementary Fig. 1. Similarly, the clone size distribution followed scaling law not in the hierarchical model, but in the NC and hNC models when one of master stem cells or one of competitive stem cells was randomly labeled (**Supplementary Fig. 2, 3**). Furthermore, when the proliferation rate of master stem cells was much higher than that of competitive stem cells, which is probably not consistent with physiological conditions, the scaling behavior collapsed (**Supplementary Fig. 4**).

**Figure 4.**
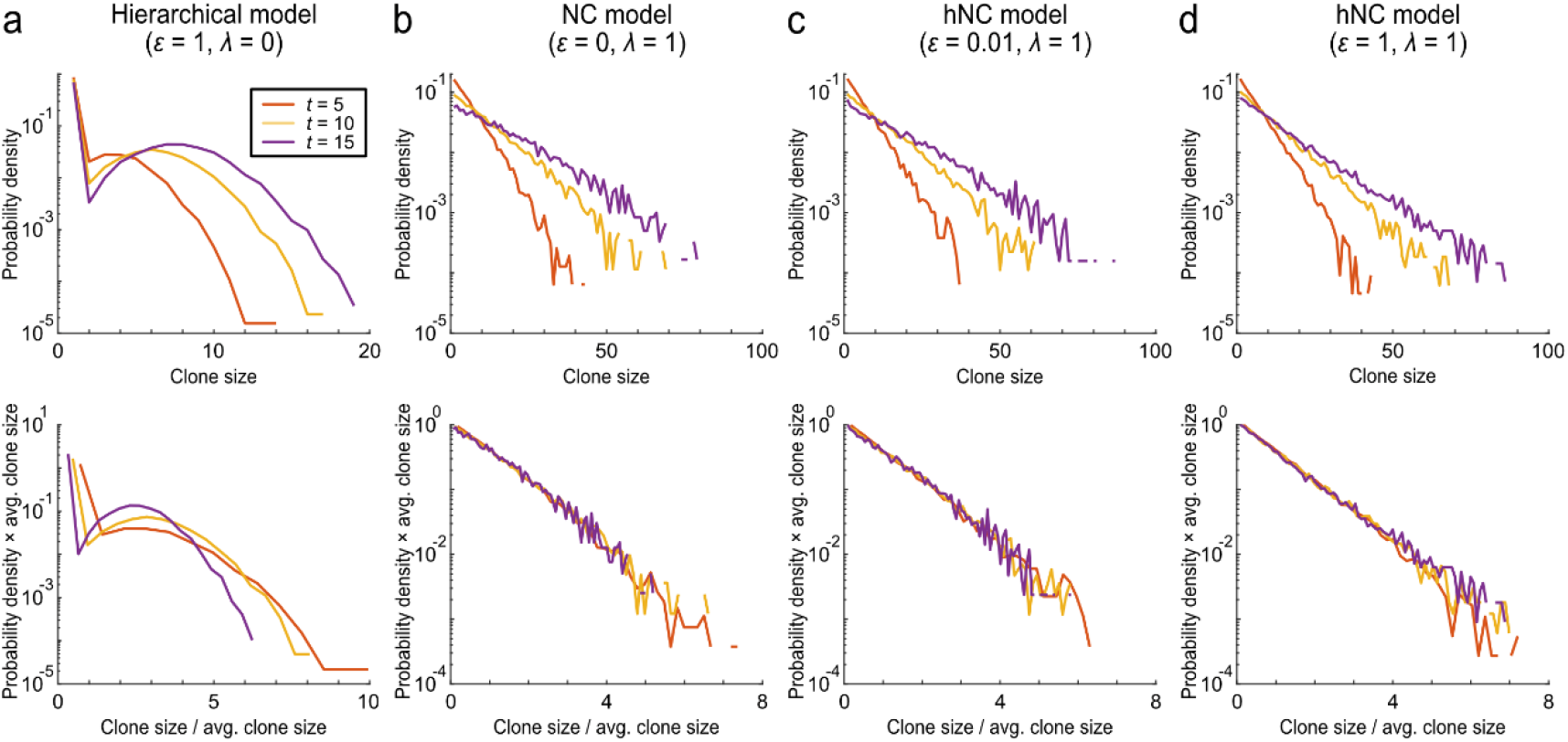
Scaling law of clone size distribution in the NC and hNC models. Probability distribution of clone size calculated by the simulation mimicking the pulse-labeling experiment. Simulations were performed 100,000 times using 10 master stem cells and 100 competitive stem cells. Cells were randomly labeled from among the master and competitive stem cells. The clone size distributions of a labeled clone were plotted at different time points in the (a) hierarchical model, (b) NC model, and (c,d) hNC model with the two conditions. The upper and lower panels show the distributions before and after scaling by the average clone size at each time point, respectively, as equation (1).

Taken together, these results indicate that the scaling law of the clone size distribution can rule out the hierarchical model and be used as an indicator of neutral competition in clonal expansion, as in the NC and hNC models. However, the scaling law does not distinguish between the NC and hNC models; therefore, it is not possible to reject the presence of master stem cells by the scaling law of the clone size distribution.

### Experimentally distinguishing the three models

Finally, we proposed a criterion for determining the mechanism of stem cell homeostasis in each tissue, that is, a criterion to distinguish the hierarchical, NC, and hNC models using experimental data (**Fig. 5a**). First, to distinguish the hierarchical model from the other two models, one needs to examine whether the scaling law of clone size distribution holds using lineage tracing techniques because we showed that the scaling law was an indicator of neutral competition, as included in the NC and hNC models (**Fig. 4**). Next, to distinguish between the NC and hNC models, one needs to examine whether clonal bursts are observed during long-term lineage tracing, as we showed that the clonal burst is unique to the hNC model (**Fig. 2**).

**Figure 5.**
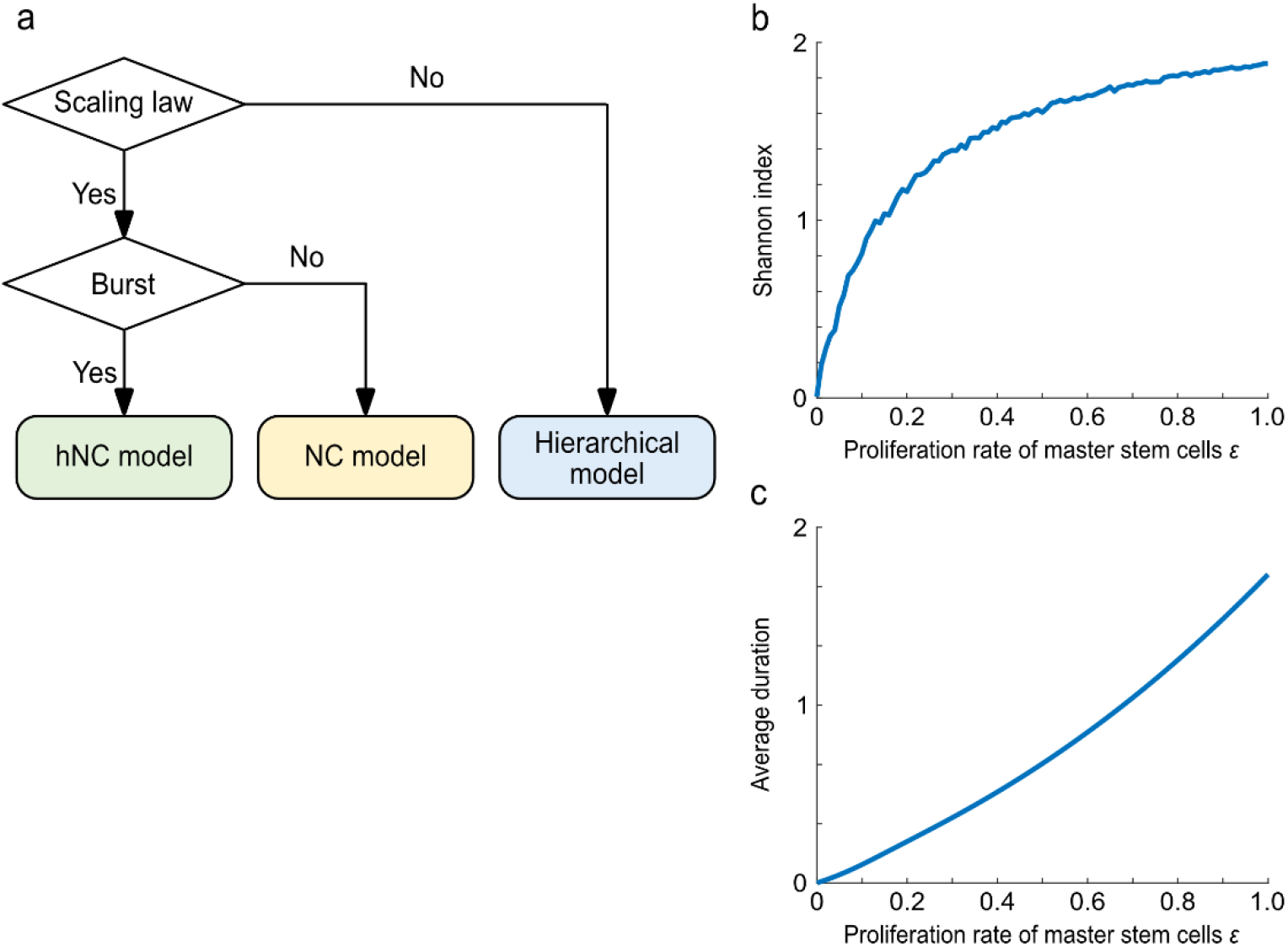
Criterion to experimentally distinguish the hierarchical, NC, and hNC models. (a) Flow chart of criterion to distinguish the hierarchical, NC, and hNC models. Scaling law of clone size distribution can be used as the indicator to distinguish the hierarchical model and other two models. Clonal bursts can be used as the indicator to distinguish the NC model and hNC model. (b, c) Two types of experimentally-measurable variables depending on the proliferation rate of master stem cells, *ε*. Shannon index representing clonal diversity (b) and average duration of all bursts (c) were plotted. *ε* can be estimated using Shannon index and/or the average duration of all bursts, both of which can be measured through lineage tracing experiments of multiple clones.

This criterion can distinguish between the three models but does not provide more detailed information on the hNC model. As shown in Fig. 2 and 3, the proliferation rate of master stem cells, *ε*, governs burst-like dynamics and thus largely affects stem cell homeostasis in each tissue. Therefore, it is important to quantify *ε* in the hNC model. Here, we propose two methods for estimating the proliferation rate of master stem cells, *ε*, in the hNC model from experimental data. The first method was based on the Shannon index *H*, which is used in ecology to measure species diversity^23^:

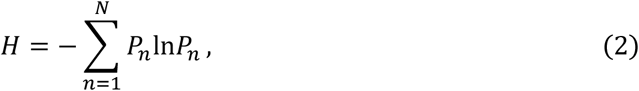

where *n* and *N* indicate the index of labeled clones and number of introduced labels, respectively, and *Pn* indicates the fraction of *n*-th labeled clonal stem cells. *Pn* can be measured by lineage tracing of multiple stem cells. Note that it is not necessary to label all stem cell clones if the total number of competitive stem cells is determined in other experiments. By simulation of our mathematical model, we showed how the Shannon index depends on *ε* (**Fig. 5b**). Based on this result, *ε* can be estimated using the Shannon index, which can be experimentally quantified by equation (2). The second method is based on the average duration of all bursts observed during lineage tracing. The average duration of all bursts of all heights increased with *ε* (**Fig. 5c**), indicating that *ε* can be estimated experimentally. When using this method, frequent and long-term lineage tracing is necessary.

## Discussion

How is stem cell homeostasis achieved in each tissue? To date, it has been controversial whether there are hierarchies among stem cells and master stem cells provide stem cells by asymmetric divisions (the hierarchical model), or whether equipotent stem cells neutrally compete with each other (the NC model) to maintain the total number of stem cells. We addressed this question by developing a mathematical model that links two existing models and represents an intermediate model named as the hNC model. Through simulation and mathematical analysis, we found unique clonal dynamics in the hNC model but not in the two existing models; the clone size repeatedly exhibited transient bursts. We also showed that the clone size distribution in the hNC model followed the scaling law, proposed as the indicator of the NC model. Based on these distinct characteristics, we proposed a criterion for discriminating the three models using experimental data.

### Experimental report supporting the hNC model

There is experimental evidence that suggests the hNC model in mammalian spermatogenesis. Kanatsu-Shinohara et al. examined the dynamics of mouse germline transmission by transplantation of spermatogonial stem cells with genetic labeling^18^. By analyzing genetic labels in offspring, they found that offspring were derived non-randomly from all labeled clonal types of spermatogonial stem cells, and periodically observed offspring derived from specific clones. This result suggests that the population size of each stem cell clone increased and decreased repeatedly, showing transient burst-like dynamics. In addition, they showed that the most undifferentiated stem cell population, that is, undifferentiated spermatogonia, was actively cycling without mitotic quiescence, whereas a more differentiated stem cell population of differentiating spermatogonia partially underwent apoptosis during differentiation.

Considering the interpretation of these findings based on the hNC model, undifferentiated spermatogonia might correspond to master stem cells that are equally cycling. In contrast, differentiating spermatogonia might correspond to competitive stem cells, because they partially undergo apoptosis, implying the presence of cell competition. Thus, in mouse spermatogenesis, it was suggested that master stem cells constantly supply competitive stem cells and competitive stem cells compete with each other, which can lead to burst-like dynamics as observed in the hNC model. Therefore, these experimental results support the hNC model, although the lineage of labeled stem cells should be directly observed to verify this model.

### Reconsideration of the hierarchical and NC model

The hNC model enabled us to reconsider the hierarchical and NC models. In the hierarchical model, there is a hierarchy among stem cells, reflected by master stem cells and non-master stem cells. In the NC model, there is no hierarchy among stem cells, which are under neutral competition. Both hierarchy and neutral competition among stem cells exist in the hNC model. Thus, even if a hierarchy is observed among stem cells and master stem cells are identified, the hNC model may be applied if cells differentiated from master stem cells compete neutrally. Similarly, even if neutral competition is observed among stem cells, the hNC model may be applied if master stem cells are present. Therefore, we may have overlooked the possibility of the hNC model by only using data from current experimental techniques.

### Comparison between the NC and hNC models

Klein and Simons theoretically suggested that the scaling law of clone size distribution can indicate the NC model^5^, and the scaling law has been confirmed by pulse-labeling of stem cells in several tissues such as the intestine^21^, epidermis^15^, and testis^24^ in mice. Although we found that the scaling law ruled out the hierarchical model, it was not able to distinguish between the NC and hNC models, because the scaling law was observed in simulation of the hNC model when master stem cells and/or competitive stem cells were pulse-labeled (**Fig. 4**; **Supplementary Fig. 2, 3**). Intuitively, the NC and hNC models show almost the same behavior in the early phase after pulse-labeling because of the small population and low proliferation of master stem cells. Thus, the scaling law of clone size distribution cannot indicate the absence of master stem cells, and it is possible that previous studies have overlooked the potential of the hNC model.

### Our criterion to distinguish the three models and stem cell markers

When studying stem cell homeostasis, it is challenging to identify distinctive characteristics of stem cells, making it difficult to define these cells in their natural context within tissues. In fact, it is difficult to determine which stem cell populations, such as master stem cells and competitive stem cells, were actually observed by a widely-used gene marker of stem cells. In this study, we proposed a criterion for distinguishing the three models by examining the presence or absence of scaling law of clone size distribution and bursts in stem cell clonal dynamics. Notably, this criterion does not depend on the strict definition of stem cells. To detect the scaling law and bursts, a minimum requirement is labeling the most undifferentiated stem cell population, which is master stem cells in the hNC and hierarchical models and competitive stem cells in the NC model. In other words, a gene marker that defines the cell population, including the most undifferentiated stem cells, is sufficient to distinguish between the three models, which means we do not need to find a gene marker specific to the most undifferentiated stem cells. It is likely to be easier to identify this population because some tissue stem cells, such as intestinal, epithelial, and spermatogenic stem cells, show distinct anatomical features. In these tissues, stem cells and differentiated cells change their physical positions in one direction during differentiation. This distinctive characteristic may enable identification of stem cell populations including the most undifferentiated cells.

## Methods

### Mathematical Model of stem cell clone dynamics

To examine stem cell clonal dynamics, we developed a mathematical model that comprehensively represents the hierarchical, NC, and hNC models. The mathematical model is based on the Moran process, which is a simple stochastic process describing population dynamics. The model is comprised of two types of stem cells: master stem cells in the closed layer and competitive stem cells in the open layer (**Fig. 1a, c, e**). The closed layer contains *K* master stem cells, which are stably maintained and not subject to competition, and provide differentiated cells named as competitive stem cells to the open layer with a proliferation rate defined as *ε*. The open layer contains *N* competitive stem cells, which are lost by their differentiation or apoptosis, and each loss is compensated by proliferation of other competitive stem cells in the open layer or master stem cells from the closed layer.

We simulated the population dynamics of distinct *K* clones of competitive stem cells. Each elementary step of the simulation involved a loss event and compensation event. For loss, one of the *N* competitive stem cells is chosen at random. The selection probability of the *k*th clone is *pk* = *nk*/*N*, where *nk* indicates the population of the *k*-th clone in the open layer. For compensation, one stem cell is chosen from the closed and open layers at random based on proliferation rate of each clone; the selection probability of the *k*-th clone is *qk* = (*nkλ* + *ε*)/(*λN* + *εK*). The total populations in the closed and open layers, that is, *K* and *N*, are constant.

To ensure time-scale consistency among the mathematical models with different parameters, the time scale was calibrated based on the average time taken per step as *t* = *m*/(*λN* + *εK*), where *m* indicates the number of steps in the simulation.

### Master equation of Moran process

To analyze clonal expansion in the Moran process, we focused on one clone of *K* clones in the open layer. The dynamics of a clone of interest can be described using the master equation (3) (**Supplementary Fig. 5**):

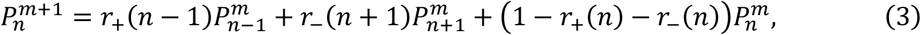

Wher 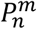 denotes the probability that the size of clones of interest is *n* in step *m*. In addition, *r*_+_(*n*) and *r*_−_(*n*) denote the transition probabilities that the clone size increases and decreases by one from *n*, respectively, as

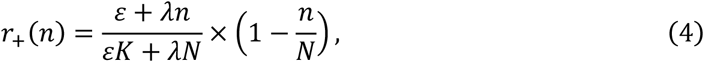

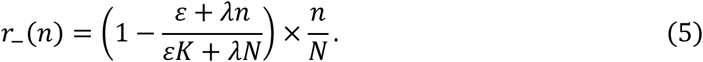

The stationary distribution of clone size 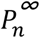 was calculated from the following detailed balance:

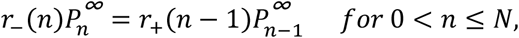

which leads to 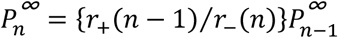. Thus,

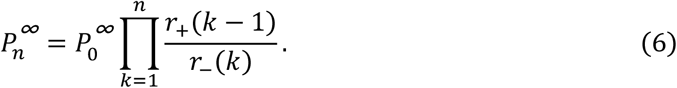

Because 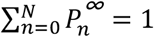, we obtained

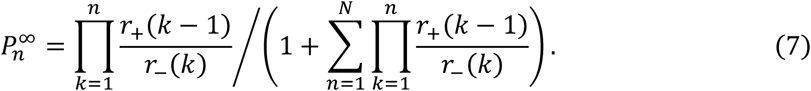

### First-passage analysis of burst-like clonal expansion

One of the distinct characteristics of the hNC model was burst-like clonal expansion (**Fig. 2c and Supplementary Fig. 1**). We analytically evaluated the generation probability and expected duration of a burst. A burst of height *H* is defined as the dynamics during which the clone size changes from 0 to *H* without returning to 0, and then returns to 0 without reaching *H* + 1 (**Fig. 3b**). Briefly, each burst generation can be separated into two processes: the forward process from 0 to *H* and backward process from *H* to 0. We evaluated this issue based on the splitting probability and first-passage time.

First, we calculated the probability of generating a burst of height *H* by multiplying the probabilities of the forward and backward processes. The splitting probability in the forward process can be described as follows:

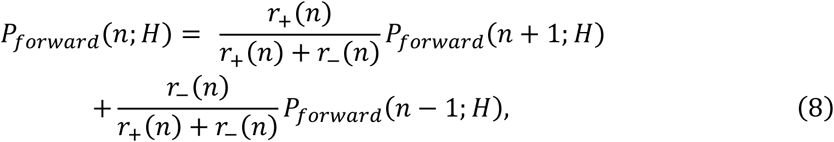

where *P*_*forward*_(*n*; *H*) indicates the splitting probability that the clone size increases from *n* to *H* without returning to 0. The boundary conditions are as follows:

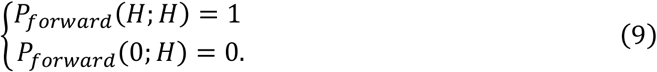

*P*_*forward*_(1; *H*) was analytically solved as

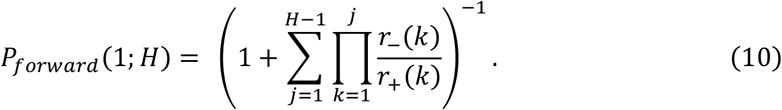

In the same manner, the splitting probability in the backward process can be described by

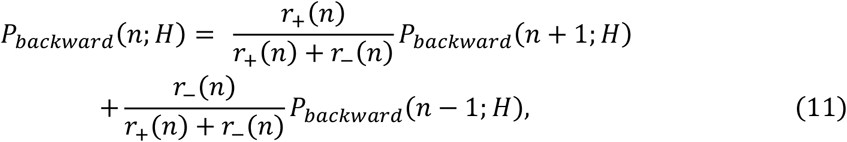

where *P*_*backward*_(*n*; *H*) indicates the splitting probability of the clone size decreasing from *n* to 0 without reaching *H*. The boundary conditions are as follows:

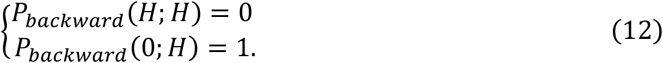

*P*_*backward*_(*H*; *H* + 1) was analytically solved as

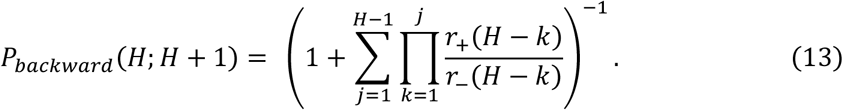

Finally, the generation probability of a burst of height *H, P*_*gen*_(*H*), was obtained using the following equation:

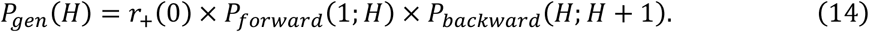

Second, we calculated the expected duration of a burst of height *H* by summing the expected durations of the forward and backward processes. The expected duration in the forward process is described by the following recurrence formula:

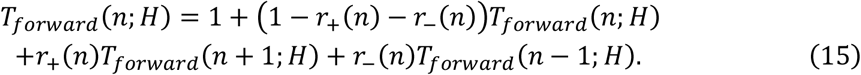

where *T*_*forward*_(*n*; *H*) indicates the expected duration in which clone size *n* increases to *H* without returning to 0 and the first term 1 indicates a unit time increment for a transition. The boundary condition at *n* = *H* is *T*_*forward*_(*n*; *H*) = 0, whereas that at *n* = 0 cannot be defined because of the absorbing boundary; therefore, this recurrence equation is intractable. To avoid this problem, we introduced the physical quantity *X*_*forward*_ defined as

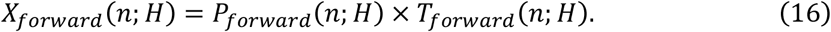

Then, the recurrence formula of *X*_*forward*_(*n*; *H*) is described by

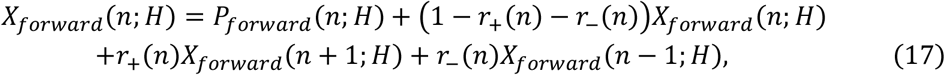

where the first term represents an increment of *X*_*forward*_(*n*; *H*), which occurs with the increment of the unit time. The boundary conditions are:

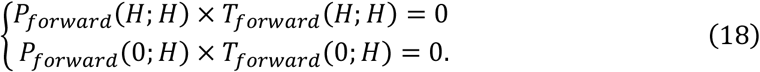

Similarly, we introduced *X*_*backward*_(*n*; *H*) = *P*_*backward*_(*n*; *H*) × *T*_*backward*_(*n*; *H*) in the backward process, where *T*_*backward*_(*n*; *H*) indicates the expected duration in which clone size *n* decreases 0 without returning to *H*. The recurrence formula of *X*_*backward*_(*n*; *H*) is described by:

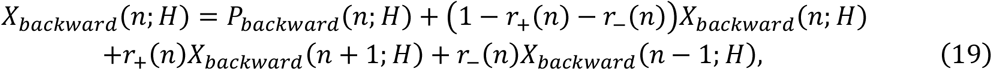

with the following boundary conditions:

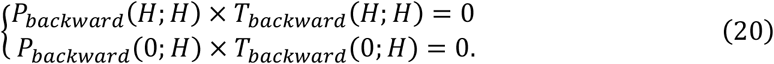

Finally, the expected duration of a burst of height *H, T*_*gen*_(*H*), was obtained using

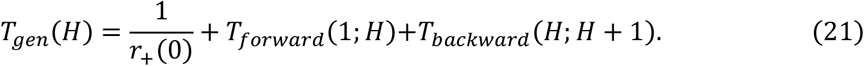

## Data availability

No datasets were generated and analyzed during the current study.

## Acknowledgements

We are grateful to Dr. Takashi Shinohara and Dr.Mito Kanatsu-Shinohara for their valuable discussions. We also thank for the research opportunity provided by the Mathematics-based Creation of Science (MACS) Program in Faculty of Science, Kyoto University (A.N. and H.N.). This study was supported in part by the Moonshot R&D–MILLENNIA Program [grant number JPMJMS2024-9] by JST, Grant-in-Aid for Transformative Research Areas (B) [grant number 21H05170], and Cooperative Study Program of Exploratory Research Center on Life and Living Systems (ExCELLS) [program number 19-102 to H.N.]. It was also supported by JSPS KAKENHI [grant number JP21J23680 to K.Y.] and Grant-in-Aid for Scientific Research (B) [grant number 21H03541 to H.N.], both from the Japan Society for the Promotion of Science (JSPS).

## Author Contributions

H.N. conceived the project. A.N., K.Y., and H.N. developed the method, A.N. implemented the model simulation and mathematical analysis, and A.N., K.Y., and H.N. wrote the manuscript.

## Figure legends

**Supplementary figure 1:**
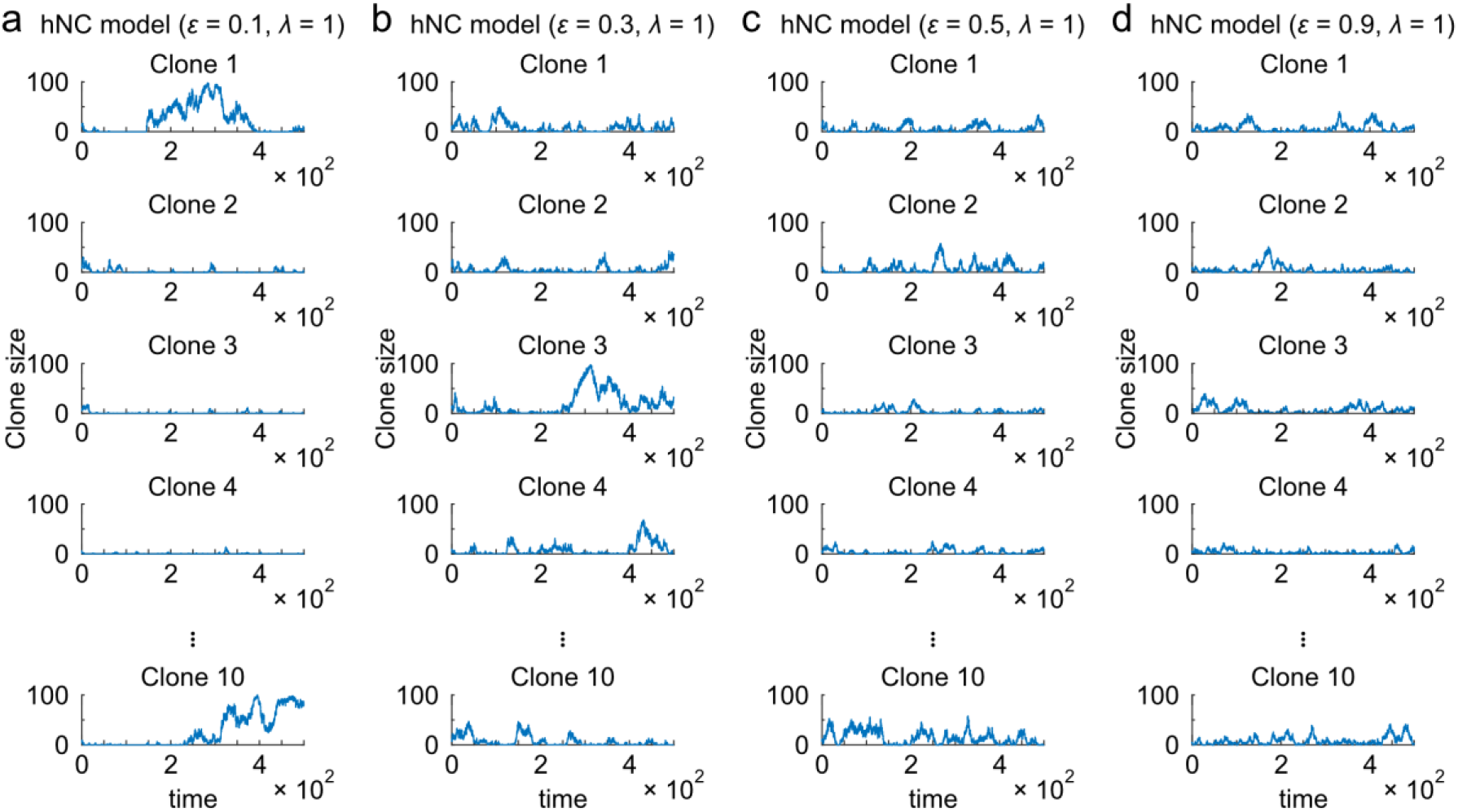
Time-series of clone sizes in the hNC model under various conditions. Time-series of clone sizes in the hNC model with different proliferation rates of master stem cells, *ε*. The simulation included 10 types of clones in 100 competitive stem cells in the open layer, in which the clone size was initially uniform, that is *n*_*k*_ = 10.

**Supplementary figure 2:**
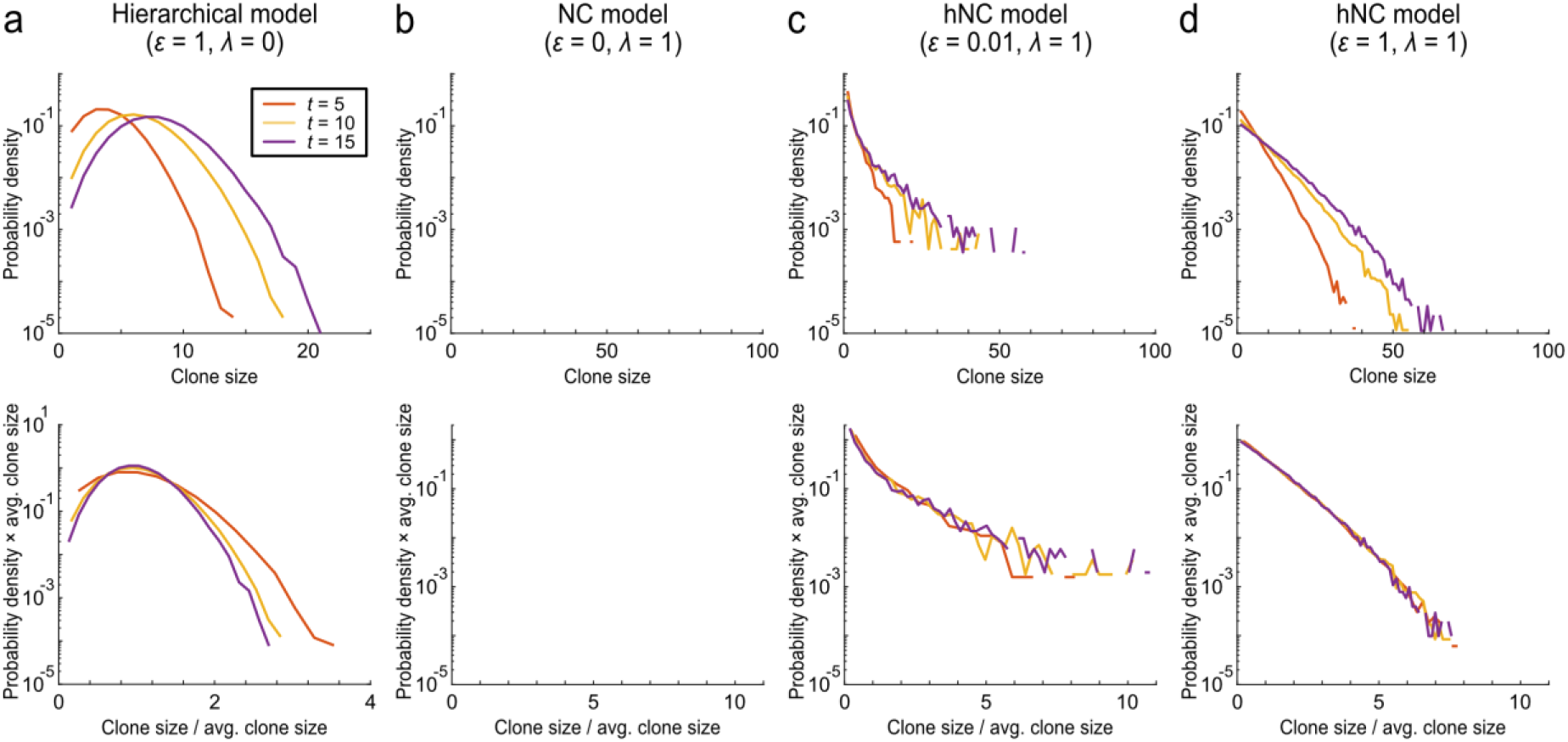
Scaling law of clone size distribution in the NC and hNC models with labelling master stem cells. Probability distribution of the population size of pulse-labeled clones. Simulations were performed 100,000 times using 10 master stem cells and 100 competitive stem cells. One of the master stem cells was randomly labeled. The labeled clonal size distributions were plotted at different time points in the (a) hierarchical model, (b) NC model, and (c, d) hNC model under the two conditions. The clone size distribution is not shown in the NC model, because the NC model has no master stem cells. The upper and lower panels show the distributions before and after scaling by the average clone size at each time point, respectively, as shown in equation (1).

**Supplementary figure 3:**
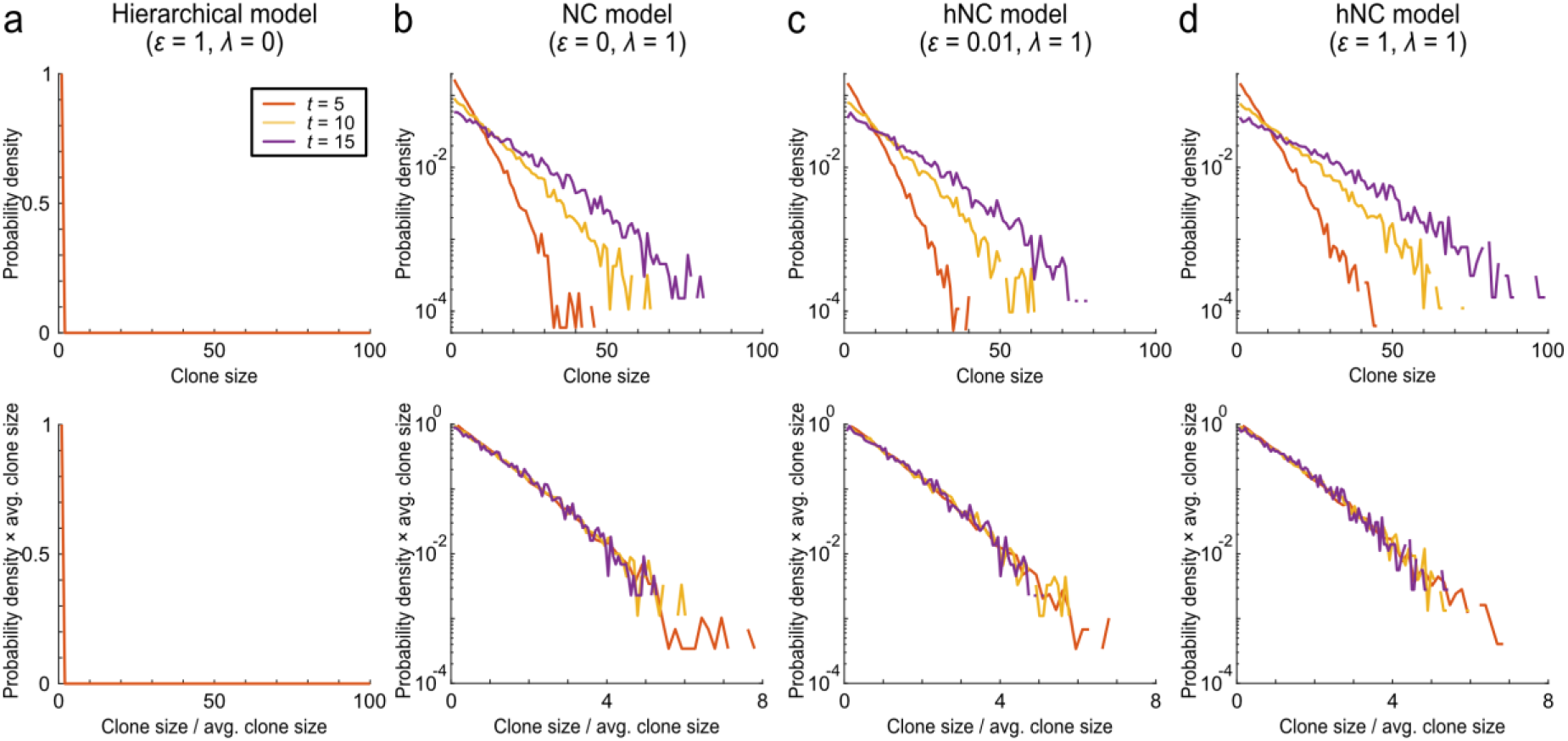
Scaling law of clone size distribution in the NC and hNC models with labeling of stem cells in the open layer. Probability distribution of the population size of pulse-labeled clones. Simulations were performed 100,000 times using 10 master stem cells and 100 competitive stem cells. One of the stem cells in the open layer was randomly labeled: non-master stem cells in the hierarchical model and competitive stem cells in the NC and hNC models. The labeled clonal size distributions were plotted at different time points in the (a) hierarchical model, (b) NC model, and (c, d) hNC model under the two conditions. Upper and lower panels show the distributions before and after scaling by the average clone size at each time point, respectively, as shown in equation (1).

**Supplementary figure 4:**
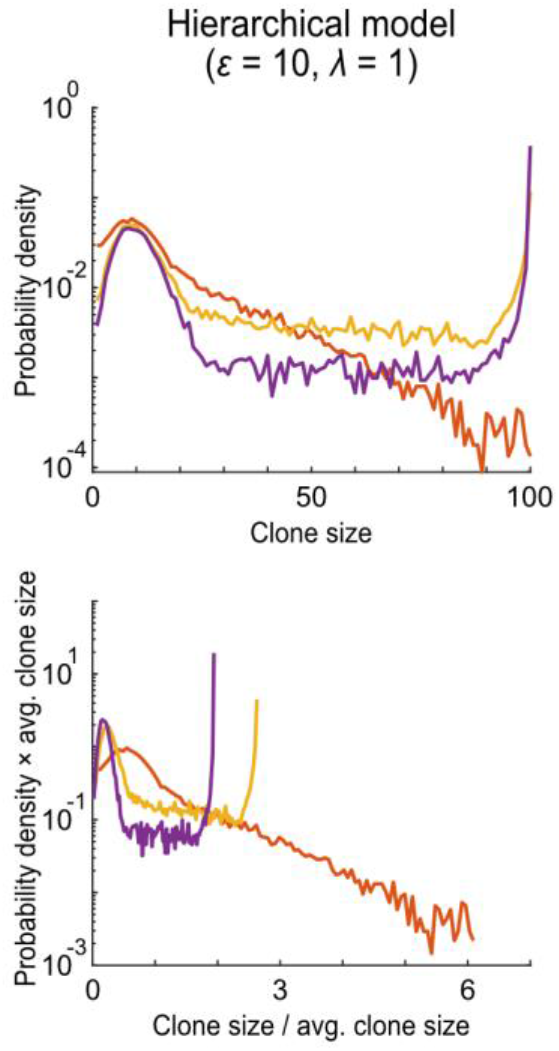
Scaling law of clone size distribution in the hNC model with high-cycling master stem cells. Probability distribution of the population size of pulse-labeled clones under the condition where the proliferation of master stem cells is much larger than that of competitive stem cells. Simulations were performed 100,000 times using 10 master stem cells and 100 competitive stem cells. One of the master and competitive stem cells was randomly labeled. The labeled clonal size distributions were plotted at different time points in the hNC model, with master stem cells being much more active than competitive stem cells. Upper and lower panels show the distributions before and after scaling, respectively, as shown in equation (1).

**Supplementary figure 5:**
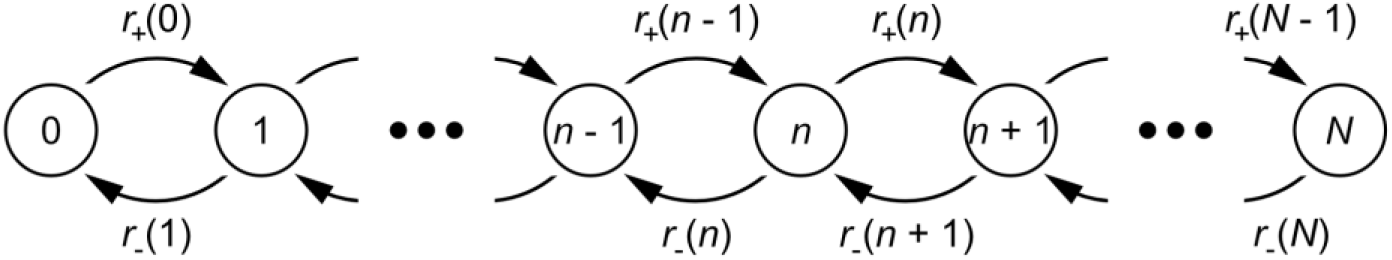
State transition diagram of clonal expansion. Clonal size stochastically changes because of neutral competition and supply from a master stem cell. *n* and *N* indicates the size of a clone of interest and the total number of competitive stem cells, respectively. *r*_+_(*n*) and *r*_−_(*n*) denote the transition probabilities that the clone size increases and decreases by one from *n*, respectively.

## References

1. Passegué E, Jamieson CHM, Ailles LE, Weissman IL. Normal and leukemic hematopoiesis: Are leukemias a stem cell disorder or a reacquisition of stem cell characteristics? Proceedings of the National Academy of Sciences. 2003;100(Suppl_1):11842–11849. doi:10.1073/pnas.2034201100

2. Ram Singh S, Burnicka-Turek O, Chauhan C, Hou SX. Spermatogonial stem cells, infertility and testicular cancer. Journal of Cellular and Molecular Medicine. 2011;15(3):468–483. doi:10.1111/j.1582-4934.2010.01242.x

3. Slaughter DP, Southwick HW, Smejkal W. “FIELD CANCERIZATION” IN ORAL STRATIFIED SQUAMOUS EPITHELIUM Clinical Implications of Multicentric Origin. Cancer. 1953;6(5):963–968.

4. Fuller MT, Spradling AC. Male and female Drosophila germline stem cells: Two versions of immortality. Science. 2007;316(5823):402–404. doi:10.1126/SCIENCE.1140861

5. Klein AM, Simons BD. Universal patterns of stem cell fate in cycling adult tissues. Development. 2011;138(15):3103–3111. doi:10.1242/DEV.060103

6. Egger B, Chell JM, Brand AH. Insights into neural stem cell biology from flies. Phil Trans R Soc B. 2008;363:39–56. doi:10.1098/rstb.2006.2011

7. Christopher SP. (No Title). Vol 7.; 1974:77–88.

8. Huckins C. The Spermatogonial Stem Cell Population in Adult Rats I. THEIR MORPHOLOGY, PROLIFERATION AND MATURATION. Vol 169.; 1971:533–558.

9. Oakberg EF. Spermatogonial Stem-Cell Renewal in the Mouse ‘. Vol 169.; 1971:515–532.

10. Simons BD, Clevers H. Strategies for Homeostatic Stem Cell Self-Renewal in Adult Tissues. Cell. 2011;145(6):851–862. doi:10.1016/J.CELL.2011.05.033

11. Cheng H, Leblond CP. Origin, differentiation and renewal of the four main epithelial cell types in the mouse small intestine V. Unitarian theory of the origin of the four epithelial cell types. American Journal of Anatomy. 1974;141(4):537–561. doi:10.1002/AJA.1001410407

12. Marques-Pereira JP, Leblond CP. Mitosis and differentiation in the stratified squamous epithelium of the rat esophagus. American Journal of Anatomy. 1965;117(1):73–87. doi:10.1002/AJA.1001170106

13. Winton DJ, Blount MA, Ponder BAJ. A clonal marker induced by mutation in mouse intestinal epithelium. Nature 1988 333:6172. 1988;333(6172):463–466. doi:10.1038/333463a0

14. Jordan CT, Lemischka IR. Clonal and systemic analysis of long-term hematopoiesis in the mouse. Genes & development. 1990;4(2):220–232. doi:10.1101/GAD.4.2.220

15. Clayton E, Doupé DP, Klein AM, Winton DJ, Simons BD, Jones PH. LETTERS A single type of progenitor cell maintains normal epidermis. Nature. 2007;446:185–189. doi:10.1038/nature05574

16. Klein AM, Nakagawa T, Ichikawa R, Yoshida S, Simons BD. Mouse Germ Line Stem Cells Undergo Rapid and Stochastic Turnover. Cell Stem Cell. 2010;7(2):214–224. doi:10.1016/J.STEM.2010.05.017

17. Snippert HJ, van der Flier LG, Sato T, et al. Intestinal Crypt Homeostasis Results from Neutral Competition between Symmetrically Dividing Lgr5 Stem Cells. Cell. 2010;143(1):134–144. doi:10.1016/J.CELL.2010.09.016

18. Kanatsu-Shinohara M, Naoki H, Shinohara T. Nonrandom Germline Transmission of Mouse Spermatogonial Stem Cells. Developmental Cell. 2016;38(3):248–261. doi:10.1016/J.DEVCEL.2016.07.011

19. Nowak M. Evolutionary Dynamics: Exploring the Equations of Life. In: Quarterly Review of Biology - QUART REV BIOL. Vol 82. ; 2007.

20. Bramson M, Griffeath D. Asymptotics for Interacting Particle Systems on Z d. Z Wahrscheinlichkeitstheorie verw Gebiete. 1980;53:183–196.

21. Lopez-Garcia C, Klein AM, Simons BD, Douglas † Winton J. Intestinal Stem Cell Replacement Follows a Pattern of Neutral Drift. Vol 330.; 2010:822–825. https://www.science.org

22. Sawyer S. A Limit Theorem for Patch Sizes in a Selectively-Neutral Migration Model. Source: Journal of Applied Probability. 1979;16(3):482–495. Accessed January 13, 2022. https://about.jstor.org/terms

23. Spellerberg IF, Fedor PJ. A Tribute to Claude Shannon (1916-2001) and a Plea for More Rigorous Use of Species Richness, Species Diversity and the “Shannon-Wiener” Index. Vol 12.; 2003. http://www.blackwellpublishing.com/journals/geb

24. Hara K, Nakagawa T, Enomoto H, et al. Mouse Spermatogenic Stem Cells Continually Interconvert between Equipotent Singly Isolated and Syncytial States. Cell Stem Cell. 2014;14(5):658–672. doi:10.1016/J.STEM.2014.01.019

